# Plastic mechanisms for unraveling a universal trade-off between water loss and respiration

**DOI:** 10.1101/2024.10.01.616113

**Authors:** Braulio A. Assis, Cameron K. Ghalambor, Eric Riddell

**Affiliations:** Department of Biology, University of North Carolina at Chapel Hill; Department of Biology, Norwegian University of Science and Technology

**Keywords:** functional conflicts, constraints, metabolic rate, respiration, plasticity, *Plethodon*

## Abstract

The expression of phenotypes is often constrained by functional conflicts between traits, and the resulting trade-offs can impose limits on phenotypic and taxonomic diversity. However, organisms can circumvent trade-offs in specific environmental contexts through phenotypic plasticity, though the mechanisms that drive trade-offs or allow organisms to resolve trade-offs are often cryptic. The trade-off between water loss and gas exchange poses a fundamental challenge for terrestrial life because the capacity to absorb oxygen is limited by the risk of desiccation from water loss across respiratory surfaces. Here, we investigated the capacity to mitigate this trade-off in a lungless salamander that breathes exclusively across its skin. We measured water loss and oxygen uptake using respirometry in field- and laboratory-based studies to identify plastic responses to environmental variation, coupled with gene expression analyses to investigate biological pathways that regulate the trade-off. Though the trade-off between oxygen uptake and water loss was generally strong, we observed that its strength changed over time in the field and laboratory. At the molecular level, we found support for antagonistic pleiotropy in multiple pathways that simultaneously affect both physiological traits (*e.g.*, via vasoconstriction and downregulation of aerobic respiration), and for mechanisms that mitigate the trade-off by affecting only one trait (*e.g.*, oxygen binding affinity, melanin synthesis). As organisms disentangle such conflicts, however, alternative trade-offs are likely to arise. Our study provides evidence that alternative pathways allow organisms to mitigate pleiotropic conflicts, which ultimately may allow greater phenotypic diversity and persistence in novel environments.

## Introduction

Ecological and evolutionary processes are universally dictated by trade-offs (Garland, 2014). Trade-offs occur because organisms are highly integrated systems such that not all biological processes can be optimized at the same time due to underlying genetic correlations, shared biological pathways, finite resources, or physical constraints (Cohen et al., 2012; Garland et al., 2022; Mauro & Ghalambor, 2020; Stearns, 1989). Though trade-offs impose limits on biological design and phenotypic expression, they also provide opportunities for organisms to express alternative combinations of phenotypes and life-history strategies across the fitness landscape (Agrawal, 2020; Meyer et al., 2015; Ricklefs & Wikelski, 2002; van Noordwijk & de Jong, 1986).

The evolutionary constraints imposed by trade-offs can either be largely fixed (*e.g.*, those governed by physical or chemical laws) or flexible (*e.g.*, those sensitive to resource availability). In turn, these differences influence the stability of trade-offs over time and across different environments (Arnold, 1992). At one end of the spectrum, fixed trade-offs are thought to impose strong evolutionary constraints on the ability of traits to evolve independently in response to natural selection (e.g. antagonistic pleiotropy; linkage), with environments selecting for opposing life-history strategies along a trade-off axis (Billerbeck et al., 2001; Conover, 1990; Ghalambor et al., 2003). On the other end, flexible trade-offs are often context dependent and can vary across individuals based on their condition (*e.g.*, the allocation of resources between competing traits; van Noordwijk & de Jong, 1986) or across changing environments (Norin & Metcalfe, 2019). In these flexible cases, variation in the strength of the trade-off can arise from environmentally driven phenotypic plasticity on the organismal scale, often achieved through physiological changes such as acclimation (Havird et al., 2020; Kelly et al., 2012; Ricklefs & Wikelski, 2002; Terblanche & Hoffmann, 2020). If traits linked by trade-offs can be regulated via phenotypic plasticity, organisms may have greater flexibility to express novel combinations of allowable phenotypes, albeit at the cost of alternative trade-offs (Sinervo & Svensson, 1988; Tikhonov et al., 2020). This combination of plasticity, integration, and conflicting functions make physiological traits chief candidates for promoting or limiting diversity and adaptation.

Water loss and gas exchange are two tightly linked physiological processes due to functional constraints inherent to terrestrial life. Both O_2_ and CO_2_ are exchanged with the atmosphere through permeable respiratory tissues that are highly vascularized, thin, and saturated with body water (Riddell et al., 2024). Being thin and saturated, respiratory tissue is also prone to experiencing high water loss rates, thus producing a fundamental trade-off between the capacity to breathe and the capacity to retain water (Tattersall, 2007; Woods & Smith, 2010). This fundamental trade-off constrains phenotypic diversity among terrestrial plants and animals (Woods & Smith, 2010), illustrating its potential to limit independent evolutionary change in these two traits (Riddell & Sears, 2020; Silvertown et al., 2015). However, there are notable evolutionary innovations that mitigate this trade-off. Nasal turbinates in birds and mammals recapture respiratory water in the nasal passage during exhalation (Geist, 2000). In some plants, CAM photosynthesis allows them to capture CO_2_ during the night when temperatures are lower, reducing costs of evapotranspiration (Heyduk, 2022). Comparative studies on amphibians have found the trade-off between gas exchange and water loss to be absent in some species and present in others (Burger et al., 2024; Riddell et al., 2024). Thus, investigating the trade-off in organisms with elevated hydric costs of respiration may reveal mechanisms through which the two traits are inherently linked or the capacity for them to be independently regulated.

The strength of a trade-off depends on the degree to which traits are independently regulated. Water loss and gas exchange tend to be linked because both processes are often regulated by the degree of blood flow to the respiratory surface (Feder & Burggren, 1985). As these traits change through plastic or evolutionary processes, changes in one trait tend to have opposing effects on the other due to shared underlying mechanisms (Riddell et al. 2024, Riddell et al., 2018; Woods & Smith, 2010). Consequently, this trade-off is likely to result in antagonistic pleiotropy as regulatory genes that increase performance of one trait might have a simultaneous reduction in performance in the other (Rose, 1982; Mauro & Ghalambor 2020). These types of gene-trait interactions likely weaken the ability of organisms to independently regulate each trait, thereby limiting the potential for independent evolutionary change. However, there are several alternative pathways that may allow organisms to regulate each trait independently. For instance, regulating oxygen binding affinity of hemoglobin may affect metabolic rate without impacting water loss (Han et al., 2011). Similarly, the synthesis of cutaneous pigments such as melanin may limit water flux across tissues (Fernandez & Koide, 2013; Ramniwas et al., 2013; Välimäki et al., 2015) without compromising gas exchange. These pathways allow for selection to favor phenotypic change in one trait independently of a related trait, thus playing a role in adaptation to drier or warmer environments.

Here, we investigated the strength of the trade-off between respiration and water loss as well as its underlying mechanisms in field-based and laboratory studies in a species of lungless salamander (*Plethodon metcalfi)*. The trade-off is particularly strong in this lineage because, being lungless, water loss and gas exchange are restricted to the same respiratory surface (Riddell et al., 2018). Specifically, *Plethodon* salamanders rely exclusively on cutaneous respiration to breathe (Gatz et al., 1974), which suggests they have a particularly limited capacity to independently regulate water loss and gas exchange. Consistent with the trade-off, laboratory experiments on lungless salamanders have demonstrated simultaneous reductions in water loss and oxygen consumption when acclimating to temperature and humidity (Riddell et al. 2018). Gene expression analyses suggest that salamanders lower water loss rates through vasoconstriction and vasoregression (Riddell et al. 2019), which provides a putative mechanism underlying the interdependence of water loss and gas uptake. However, the mechanisms associated with both traits simultaneously or with each trait independently have yet to be explored.

We evaluated the strength and consistency of the trade-off in *P. metcalfi* throughout their active season in individuals recently collected from nature to understand whether salamanders are capable of mitigating the trade-off. We also combined acclimation experiments in the laboratory with gene expression analysis to identify gene-trait associations underlying the independent and dependent regulation of water loss and gas uptake. We specifically investigated genes or gene ontologies associated with pathways related to processes that would influence both oxygen uptake and water loss (*i.e.*, pathways consistent with the trade-off) or with one trait but not the other (*i.e.*, independent regulation of water loss or gas uptake). Thus, our study reveals whether organisms adjust phenotypes along axes that are consistent and inconsistent with trade-offs, which ultimately contributes to generating phenotypic and taxonomic diversity in nature.

## Methods

This study is divided into two components (*seasonal acclimatization* and *laboratory acclimation*) that measured oxygen uptake and skin resistance to water loss in response to changing environmental conditions. The “*seasonal acclimatization study*” investigates variation in the linkage between oxygen uptake and skin resistance to water loss in salamanders recently collected from nature throughout their active season (May to October), and thus identifies the strength of the trade-off as salamanders respond to environmental change in nature. The “*laboratory acclimation experiment*” investigates plasticity in oxygen uptake and skin resistance in response to controlled temperature and humidity in the laboratory, combined with gene expression analyses on the skin. Thus, the laboratory experiment helps to identify how these traits are linked from an organismal and mechanistic perspective while minimizing variation due to the environmental history experienced in nature. In both studies, we used the volume of oxygen uptake (*V*O_2_) as a metric of oxygen uptake, and skin resistance to water loss (*r_i_*) as the metric of water loss. Skin resistance to water loss is defined as the ratio between the water vapor density gradient and the cutaneous water loss rate by surface area of the organism (Riddell et al., 2017; Spotila & Berman, 1976). By including the gradient controls for the evaporative demand of the air, we are able to isolate the physiological changes in water loss.

### Seasonal acclimatization study

We analyzed oxygen uptake and water loss rates in 181 salamanders divided into five time points between May and September 2015 (May, early July, late July, August, and September). Animals were collected from the Balsam Mountain Range in the Nantahala National Forest (35° 20′ N, 83° 4′ W) across an elevational gradient spanning 1200m and 1600m. Collection occurred in sites randomly generated on QGIS v2.1 and at least 100m away from roads. Conditions were variable throughout this period, with highest temperatures occurring in late July, and highest humidity occurring in August (Riddell et al., 2018). Collection of oxygen uptake and water loss data in the laboratory is described below (*Respirometry trials*). To ensure that salamanders were in a postabsorptive state, we maintained them at 15°C and in the absence of food for one week prior to respirometry trials. We also measured temperature and humidity throughout the season to understand the conditions salamanders experienced. We randomly generated 15 coordinates within the collection sites to deploy temperate-humidity loggers (Hygrochrons; Maxim Integrated), which were placed on the surface of the forest floor in a hardwire mesh frame to protect the logger from moving. Data were collected every 20 minutes (*n* = 2,939). At each time point, we report the temperatures (°C) and vapor pressure deficits (VPD; kPa), which describes the difference between the amount of moisture in the air and the amount it can hold when saturated. To calculate VPD from humidity data, we used the equations described in Riddell *et al*. (2018).

### Laboratory acclimation experiment

In May 2016, we collected 120 salamanders to investigate the potential for plasticity in the trade-off between oxygen uptake and skin resistance to water loss in controlled laboratory conditions. For salamanders to become accustomed to laboratory conditions, we kept them for one month in individual containers (17cm x 17cm x 12cm) with moist paper towels. Salamanders were maintained in an incubator (Percival, Inc.; Model #I-36VL) that cycled through a temperature regime that ranged between 8.5°C and 15°C, mimicking temperatures they experience above and below ground. After this period, we collected baseline measurements of oxygen uptake and skin resistance to water loss using respirometry trials (see below). We then evenly distributed the 120 individuals across four experimental groups consisting of temperature and humidity exposures in a 2x2 factorial design (temperature cycles ranging either between 8.5°C and 15°C, or between 15°C and 22.5°C; vapor pressure deficits of either 0.2 or 0.4 kPa).

On each night during the acclimation experiment, we moved salamanders into activity enclosures (17cm x 17cm x 12cm) to ensure salamanders experienced ambient humidity in the incubator. The enclosures consisted of dry soil as substrate and a hardwire mesh roof to allow air from the incubator to circulate. Exposures in activity enclosures happened for a period of three hours between 2100 and 0600 hours for five consecutive nights per week (followed by one night of respirometry trials and one night of rest) for four consecutive weeks. Each individual was weighed to the nearest 0.001 g before and after each bout of simulated activity. We monitored the body mass lost after each exposure period to ensure no animal lost more than 10% of their baseline mass, in which case that individual would not participate in another exposure period until recovery. This outcome occurred in 1.1% of all exposures. To maintain all salamanders in a postabsorptive state, they were not fed during the acclimation experiment. At the end of the acclimation experiment, we euthanized each individual for gene expression analysis and selected 8 individuals from each of the four experimental groups (*n* = 32) for sequencing. Because the skin is the tissue directly involved in cutaneous respiration and resistance to water loss, we dissected a portion of the dorsal skin for total RNA extraction (see *RNA sequencing*).

### Respirometry trials

We used the same respirometry system for the seasonal acclimatization and laboratory acclimation studies. We measured oxygen consumption and skin resistance to water loss using a flow-through respirometry system (Sable Systems Int. [SSI], Las Vegas, NV). In our system, temperature was controlled using a Percival incubator and maintained at 18°C while a subsample pump (SS4; SSI) constantly pumped air through a bubbler bottle which saturated the airstream. This saturated airstream then continued through a dew point generator (DG4; SSI) that controlled the water vapor content (vapor pressure deficit = 0.5 kPa). Then, the airstream was separated using a flow manifold (SSI) that also allowed adjustment of flow rates (180 mL/min). These adjusted airstreams were then passed through acrylic chambers (16 × 3.5 cm; volume c. 153 mL) containing the salamander, suspended over hardwire mesh. This chamber design was implemented to promote stereotypical posture during activity and to minimize the potential for posture to confound water loss rates. Although air constantly flowed through all chambers, data collection was cycled through one chamber at a time using a multiplexer (MUX8; SSI). Out of a given chamber, the airstream passed through a water vapor analyzer (RH300; SSI) and lastly through a flow meter and a dual differential oxygen analyzer (Oxzilla; SSI). Conversion of raw data outputs from the respirometry system into physiological traits is described in Riddell *et al*. (2018). Both studies were approved by the Institute for Animal Care and Use Committee (IACUC) at Clemson University (#2014-024) with approvals from the North Carolina Wildlife Resource Commission (#16-SC00746), the U.S. Fish and Wildlife Service (#16-SC00746), and Nantahala National Forest.

### Statistical analyses

We determined the strength, consistency, and changes in the trade-off using linear regressions with oxygen uptake (*V*O_2_) as a function of resistance to water loss (*r*_i_). For the seasonal acclimatization study, we analyzed the relationship between *V*O_2_ and *r*_i_ for each time point separately due high multicollinearity between time point and resistance to water loss (as determined by generalized variance inflation factor (GVIF) > 100, Fox & Monette, 1992). Each analysis also included body mass as a covariate to account for the effects of body size on oxygen uptake (Hayes, 2001).

For the laboratory acclimation experiment, we assessed the strength of the trade-off before and after laboratory acclimation, and plasticity in the trade-off over the course of the experiment (*n* = 120). For the first model, we regressed initial *V*O_2_ on initial *r*_i_ before acclimation, and for the second model, we regressed final *V*O_2_ on final *r*_i_ after the experimental acclimation. Lastly, we regressed the change in *V*O_2_ (*V*O_2_) on the changes in *r*_i_ (Δ*r*_i_) after the experimental acclimation. We selected *r_i_* as the independent variable as changes in resistance to water loss are suspected to cause changes in gas exchange, but not vice versa. For all models, body mass was included as a covariate. We also included interactions between treatments (temperature x humidity) and skin resistance for the second and third models to test if the strength of the trade-off changed in response to the experimental treatments. Due to the lack of an effect of the treatments and high collinearity between treatments and skin resistance (see *Supplementary materials*, table S1), we report results from the simplified model (without the interactions with treatment).

### RNA sequencing

We extracted total RNA from the dorsal skin of 32 salamanders immediately after the final respirometry measurements in the laboratory acclimation study. We euthanized the salamanders by immersion in liquid nitrogen and removed dorsal skin tissue using a flame-sterilized razor blade. We then immersed the tissue samples in TRIzol reagent (Item#: 15596026; Thermo Fisher Scientifics, 168 Third Avenue, Waltham, MA 02451), which was then homogenized using a motorized tissue homogenizer with sterilized pestle (Item#: UX-44468-25; Argos Technologies Inc., 625 E. Bunker Ct., Vernon Hills, IL 60061). Tissue homogenates were stored in -80°C until further downstream processing.

To purify samples for library preparation, we used the RNeasy Mini Kit for total RNA (ID: 74136; Qiagen®, 1001 Marshall St., Redwood City, CA 94063), and we reduced DNA contamination by treating the samples with DNase. We quantified total RNA and assessed integrity using a Qubit Fluorometer (Thermo Fisher Scientific) to ensure sufficient RNA of high quality in each sample and prepared libraries using the Illumina TruSeq® mRNA Stranded Kit (Product#: RS-122-2101, Illumina, Inc., 5200 Illumina Way, San Diego, CA 92122). We then used the Qubit Fluorometer to quantify the amount of cDNA in each sample prior to sequencing.

We sequenced single-end 100-bp reads across four lanes of an Illumina HiSeq2500 platform at the Genomics Facility at Cornell University. To avoid lane biases, individuals from each treatment were distributed randomly and evenly across the four sequencing lanes. After sequencing, we trimmed sequencing adapters from the raw reads using Trimmomatic (v. 0.36) and evaluated error probabilities for all reads using ConDeTri (v. 2.3), which removed low-quality bases based on default parameters. Afterwards, we used FastQC (version 0.11.5) to ensure that quality scores were > 30 for all sequences. The *P. metcalfi* transcriptome assembly is described in Riddell *et al*. 2019 and was based on 48 tissue samples (36 skin and 12 heart samples).

### Gene-trait associations

We used Pearson’s correlations to investigate transcriptomic associations with physiological traits using two thresholds. We used a higher correlation threshold (99^th^ quantile of absolute correlations) to evaluate individual genes correlated with each or both traits, and we used a lower threshold (90^th^ quantile of absolute correlations) to investigate functional enrichments on larger sets of genes. For genes strongly aligned with the trade-off, we selected genes that were above the 99^th^ quantile of absolute correlations both with *ΔVO*_2_ (|r|*_ΔVO_*_2_ > *Q*_99_) and *Δr*_i_ (|r|*_Δr_*_i_ > *Q*_99_). We evaluated genes in this set individually for potential associations with processes that might affect both water loss and oxygen uptake. We also tested for antagonistic pleiotropy by assessing the sign of the relationship between the gene and trait of interest.

Evidence of negative pleiotropy would be indicated by a positive relationship between the expression of a gene on one trait but a negative relationship for the other trait. Next, we selected a larger set of genes for a GO enrichment analysis. These were genes above the 90^th^ quantile of *ΔVO*_2_ & *Δr*_i_ (|r|*_ΔVO_*_2_ > *Q*_90_ & |r|*_Δr_*_i_ > *Q*_90_).

To identify putative pathways underlying the independent regulation of skin resistance and gas uptake, we determined which genes were correlated with changes in one physiological trait while being uncorrelated to changes in the other trait. We investigated individual genes that were above the 99^th^ quantile of absolute correlations with *r*_i_ (|r|*_Δr_*_i_ > *Q*_99_) and below the Q_90_ for |r|_ΔVO2_ and, reciprocally, genes with |r|_ΔVO2_ > Q_99_ and |r|_Δri_ < Q_90_. Lastly, we used a lower threshold to obtain a larger set of putative genes for a GO enrichment analysis. These sets were genes with |r|_Δri_ > Q_90_ & |r|_ΔVO2_ < Q_90_ and |r|_ΔVO2_ > Q_90_ & |r|_Δri_ < Q_90_. These analyses assume a continuous relationship between transcript abundance and phenotypic expression. However, threshold effects induced by low-level or discrete changes in gene expression on phenotypes (Reid & Acker, 2022) may not be captured by this approach.

### Weighted gene co-expression network analysis

Gene co-expression networks can reveal functional relationships among genes that reflect known biological pathways and networks (Campbell-Staton et al., 2018). We used a weighted gene co-expression network analysis (WGCNA) to identify networks of genes with correlated expressions that are then assigned into modules of co-expression (Zhang & Horvath, 2005). These modules allow us to assess whether candidate genes (identified in the correlation-based analysis above) are controlled by a shared mechanism (*i.e.*, if they belong to the same co-expression module) or expressed independently (*i.e.*, if they belong to many different modules). If candidate genes are assigned to different modules, we can infer that the trait is regulated by multiple independent pathways. Conversely, candidate genes in the same module would suggest the trait is potentially regulated as part of an integrated pathway. To provide an additional (but independent) analysis relating gene expression to phenotypic variation, we used a principal components analysis (PCA) to summarize individual variation in phenotypic change in *V*O_2_ and *r*_i_, and then assessed the correlation between the first two principal components and the eigenvalues of the WGCNA modules. In this analysis, the first principal component (PC1) encompasses the variation consistent with the trade-off (the correlation between the change in *V*O_2_ and *r*_i_). The second principal component encompasses the variation orthogonal to the trade-off and thus describes the independent changes in both *V*O_2_ and *r*_i_. We note that PC2 is not capable of determining which genes are associated with the independent regulation of *V*O_2_ or *r*_i_ (as done in the correlation analysis above), as they are both integrated into the same principal component.

After removing all genes with less than 10 counts across 28 samples, we retained 9346 genes for the WGCNA. We then normalized all gene counts using the *varianceStabilizingTransformation* function in DESeq2 and removed any genes with missing data or variance of zero. After these steps, one individual from the cool/wet treatment failed quality control and was removed from downstream analyses. The gene co-expression networks are based on pairwise Pearson’s correlations using the *blockwiseModules* function and a soft thresholding of 4. We defined a gene network dendrogram based on average linkage hierarchical clustering along with a gene dissimilarity matrix. Afterwards, we used the Dynamic Tree Cut method to merge modules with high correlation (height-cut = 0.25).

### Gene Ontology enrichment analyses

To identify functional enrichments associated with sets of genes, we used the R package *goseq* (Young et al., 2010) for Gene Ontology (GO) enrichment analyses. We excluded GO terms associated with fewer than 10 and more than 500 genes to reduce enrichment sensitivity and redundancy. To reduce positive sequencing bias for longer transcripts, transcript length for all genes was integrated into the analysis. The entire set of genes in the *P. metcalfi* skin transcriptome consisted of 9,346 unique transcripts. However, we excluded any uncharacterized transcripts from the analysis, resulting in a background set of 6,953 genes.

## Results

### Water loss/oxygen uptake trade-off

In four of the five time points between May and September, salamanders in the seasonal acclimatization study exhibited a significant trade-off between skin resistance (*r_i_*) and oxygen uptake (*V*O_2_; Fig. 1), indicated by a negative relationship between the traits. Each analysis also accounted for body mass, which was significantly related to oxygen uptake in each model (p < 0.05 in each analysis). However, salamanders collected in August did not show a significant relationship between the two traits (β = -0.002, p = 0.97). August was one of the warmest and wettest months of the year (Supplementary Figure S2). Though the trade-off seems mostly consistent throughout their active season, the lack of a trade-off in August suggests the two traits can be regulated independently under certain conditions.

**Figure 1:**
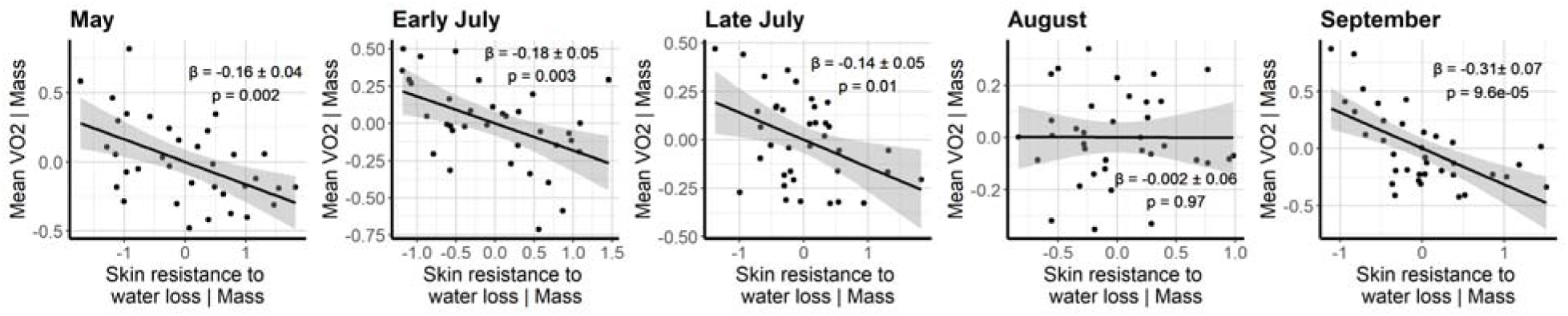
Fluctuating strength of the trade-off between skin resistance and water loss throughout the active season in *P. metcalfi*. During August, no significant association between the two traits was observed. Plots are partial regressions that account for individual body mass.

Similarly in the laboratory acclimation study, the trade-off between skin resistance and oxygen uptake was significant in individuals prior to acclimation (Fig. 2A), but not after (Fig. 2B). We also found evidence for a change in the trade-off in the plasticity between traits, with increases in skin resistance (Δ*r*_i_) being associated with reductions in oxygen uptake (Δ*V*O_2_) over the course of the experiment (Fig. 2C; p < 0.001). These results indicate that the trade-off between the two traits constrains acclimation responses. The results also show that the strength of the trade-off also changes in response to acclimation. The temperature and humidity treatments had no significant effect on the trade-off or its change during the laboratory experiment (see *Supplementary materials*, table S1). We also note that the sequenced individuals (*n* = 32) exhibited the same patterns as all individuals in the acclimation experiment (*n* = 120, Supplementary Figure S3).

**Figure 2:**
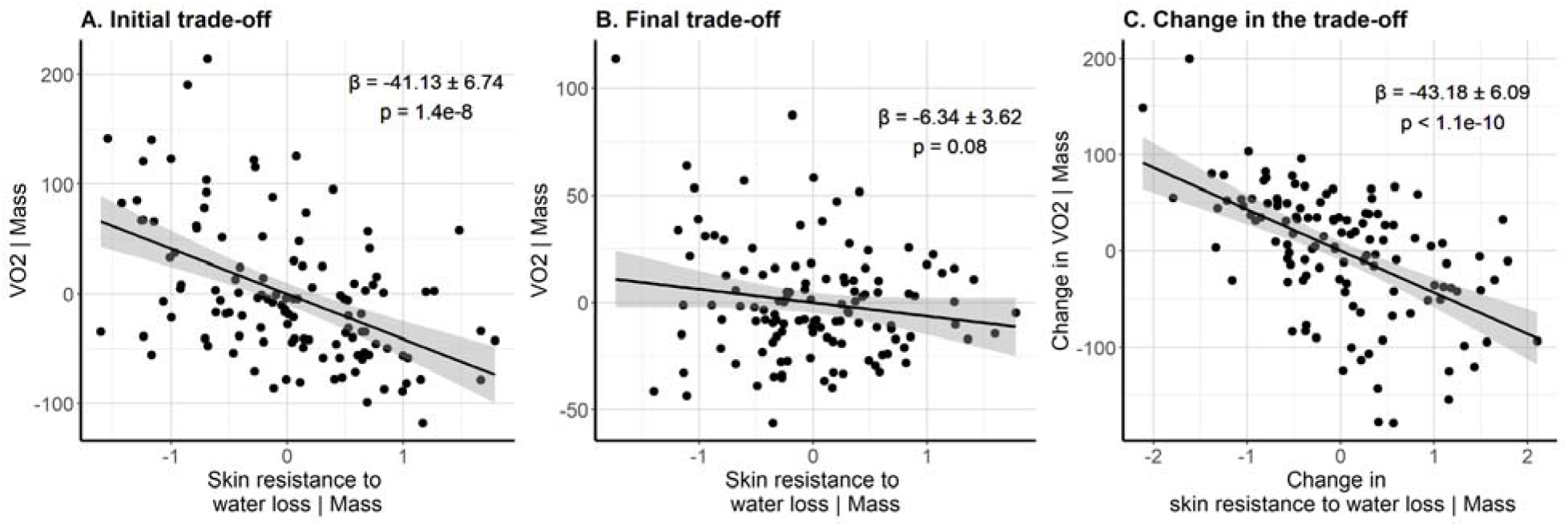
Partial residual plots accounting for body mass for the oxygen uptake/water loss trade-off during the laboratory acclimation experiment. A. Skin resistance and oxygen uptake had a significant negative association at the beginning of the experiment. B. After the acclimation experiment, the relationship between the two traits was marginally non-significant. C. Increases in skin resistance were strongly associated with decreases in oxygen consumption after the acclimation experiment.

### Gene expression associated with the trade-off

The intersection between |r|_Δri_ > Q_99_ and |r|_ΔVO2_ > Q_99_ was represented by 19 genes, 10 of which are characterized (Supplementary Table S4). Consistent with the underlying functional basis of the trade-off, oxygen uptake and resistance to water loss were inversely correlated with the expression of each gene (Figure 3A, 3B), consistent with evidence for antagonistic pleiotropy. The highest combined absolute correlation with both traits was for pyruvate dehydrogenase isozyme 4, which is associated with the downregulation of aerobic respiration and glucose metabolism (Pettersen et al., 2019). In the principal component analysis, PC1 explained 77.3% of the variation in the physiological traits (*i.e.*, change in the two traits that is consistent with the trade-off). The WGCNA found differentially expressed genes were distributed across 27 modules. The WGCNA also indicated that one co-expression module was significantly associated with PC1 (*magenta*; r = -0.37, p = 0.037). The gene ontology analysis of the *magenta* module (203 genes) resulted in 24 enriched terms (Supplementary table S5), the most significant of which was *negative regulation of angiogenesis* (3 overexpressed genes). Of the 19 genes most correlated with the trade-off, three were included in the magenta module (one of which is characterized).

**Figure 3:**
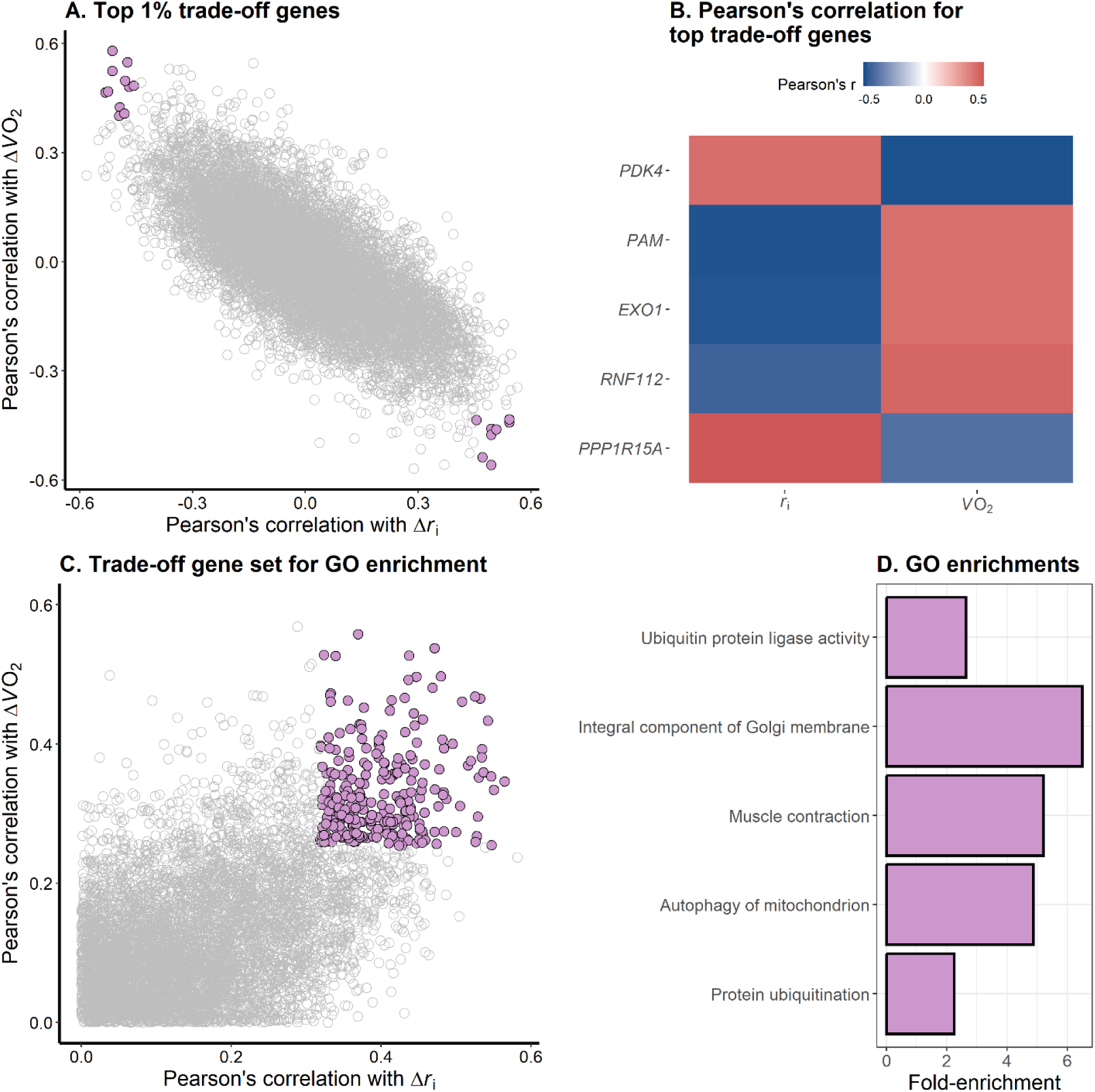
Genes and processes putatively involved in the trade-off between oxygen uptake and water loss A. Pearson’s correlations with changes in skin resistance to water loss (Δr_i_) and with changes in oxygen uptake (Δ*V*O_2_) throughout the laboratory acclimation experiment for 9,346 transcripts sequenced from *P. metcalfi* skin tissues. Highlighted points correspond to genes above the 99^th^ percentile of absolute correlations with both Δr_i_ and Δ*V*O_2_ (n=19). B. Heatmap for the genes with the highest combined correlation across Δ*r*_i_ and Δ*V*O_2_. C. Set of genes for functional enrichment analysis. Highlighted points correspond to genes above the 90^th^ percentile of absolute correlations with both Δ*r*_i_ and Δ*V*O_2_ (n=267) out of 6,953 characterized transcripts in the *P. metcalfi* skin transcriptome. D. Five most significant functional enrichments for the set of genes in panel C.

**Figure 4:**
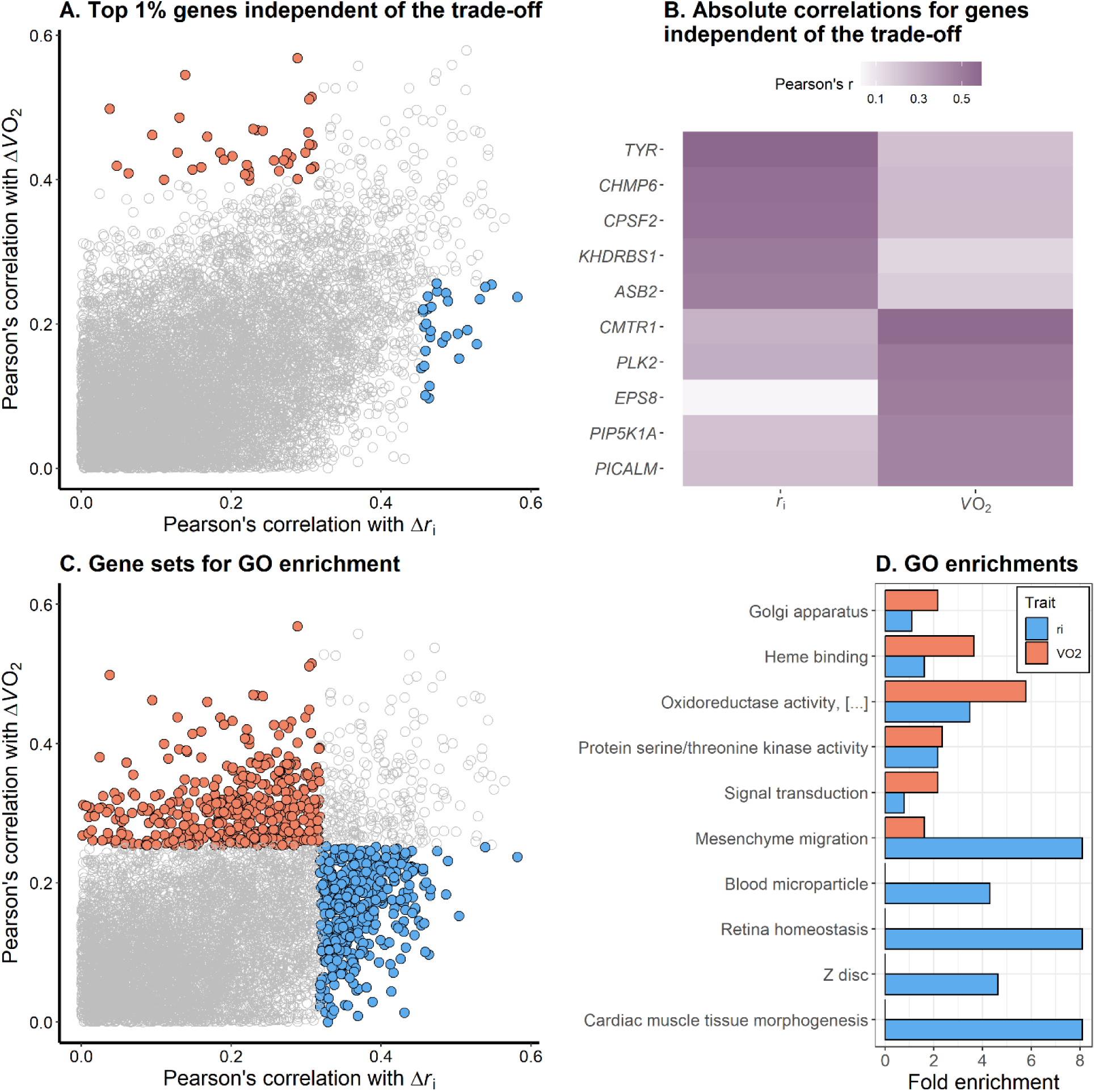
Genes and processes associated with the independent regulation of oxygen uptake and skin resistance to water loss. A. Absolute values for Pearson’s correlations with changes in skin resistance to water loss (Δ*r*_i_) and with changes in oxygen uptake (Δ*V*O_2_) throughout the laboratory acclimation experiment for 9,346 transcripts sequenced from *P. metcalfi* skin tissues. Points in blue correspond to genes above the 99^th^ percentile of absolute correlations with Δ*r*_i_ and below the 90^th^ percentile of absolute correlations with Δ*V*O_2_ (n=30), while points in red correspond to the reciprocal set for Δ*V*O_2_ (n=38). B. Heatmap for the five genes with the highest correlations for each of the two sets highlighted in A. C. Set of genes for functional enrichment analysis. Points in blue correspond to genes above the 90^th^ percentile of absolute correlations with Δ*r*_i_ and below the 90^th^ percentile of absolute correlations with Δ*V*O_2_ (n=429) out of 6,953 characterized transcripts in the *P. metcalfi* skin transcriptome, while points in red correspond to the reciprocal set for Δ*V*O_2_ (n=429). D. Five most significant functional enrichments for each set of genes in panel C.

Next, we investigated functional enrichments in the larger set of genes correlated with both traits (|r|_Δri_ > Q_90_ and |r|_ΔVO2_ > Q_90_; n=267; Supplementary table S6). Among the five most significant enrichments, at least three terms were associated with protein degradation and downregulation of aerobic metabolism (*Ubiquitin protein ligase activity*, 7 genes; *autophagy of mitochondrion*, 3 genes; *protein ubiquitination*, 7 genes). The third most significant enrichment was for *muscle contraction* (3 genes). Further, 8.9% of genes in this set (24 out of 267) also occurred within the *magenta* module.

### Genes associated with the independent regulation of physiological traits

Thirty genes had transcription levels with the highest correlations with Δ*r*_i_ (|r|_Δ*r*i_ > Q99) while being uncorrelated with Δ*V*O_2_ (|r|_Δ*V*O2_ > Q99); Supplementary table S7). The strongest correlation with Δ*r*_i_ was with the expression of tyrosinase, *TYR* (|r| = 0.58, p = 0.001), which is a precursor to melanin synthesis. Among functional enrichments associated with the larger set of 429 genes (|r|_Δri_ > Q_90_ & |r|_ΔVO2_ < Q_90)_; Supplementary table S8), the most significant was for the biological process *mesenchyme migration* (5 genes). Further, the cellular component *melanosome* was significant (4 genes, p = 0.02). Thirty-eight transcripts had the highest correlations with Δ*V*O_2_ (|r|_Δ*V*O2_ > Q99) *V*Owhile being Δ*r*_i_ (|r|_Δ*r*i_ > Q99); Supplementary table S9). The highest correlations with Δ*V*O_2_ were for *CMTR1* (|r| = 0.57, p < 0.001), associated with RNA cap formation (Liang et al., 2023), and *PLK2* (|r| = 0.51, p = 0.003), which regulates the mitotic spindle orientation (Villegas et al., 2014). For the larger set of 429 transcripts (|r|_Δvo2_ > Q_90_ & |r|_Δri_ < Q_90_; Supplementary table S10), (|r|_ΔVO2_ > Q_90_ & |r|_Δri_ < Q_90_) the second most significant enrichment was for the term *heme binding* (9 genes).

In the principal component analysis, PC2 explained 22.7% of the variation (*i.e.*, phenotypic change orthogonal to the trade-off). One WGCNA module was significantly correlated with PC2 (*darkorange*, r = -0.36, p = 0.043). The two most significant enrichments in the GO analysis of the *darkorange* module were *immune system process* and *immune response*, with several other enriched terms associated with the regulation of unspecific cellular processes (Supplementary table S11). None of the genes most correlated with the independent regulation of gas uptake and skin resistance were included in the *darkorange* module. Among the broader set of genes, 1.2% (5 out of 429) occurred within the *darkorange* module.

## Discussion

Functional conflicts between traits can constrain phenotypic diversity and evolutionary change by producing trade-offs (Arnold, 1992; Gupta et al., 2022; Martin, 2014; Rosenfeld et al., 2020; Wooliver et al., 2016). However, the degree to which organisms may leverage pathways that mitigate a trade-off by allowing traits to occupy new phenotypic spaces and circumvent conflicts remains unclear (Hashemi et al., 2024; Shoval et al., 2012). In this study, we demonstrate that the fundamental linkage between cutaneous gas exchange and water loss in a lungless salamander varies in strength through time (Fig. 1, 2), which suggests that salamanders have the capacity to regulate functional conflicts related to this trade-off (Buehler et al., 2012). Similarly, we identified genes associated with the independent regulation of both gas uptake and skin resistance to water loss. In contrast to these independent effects, we also identified generally strong associations between gas uptake and skin resistance in the seasonal and acclimation studies as well as multiple genes expression patterns consistent with antagonistic pleiotropy (Fig. 3), highlighting the strong interdependence of the two traits. Therefore, patterns of gene expression can not only identify the putative mechanisms that drive the trade-off, but also the mechanisms that mitigate the trade-off. These findings provide new insights into the mechanisms through which organisms can regulate trade-offs, the environmental contexts in which this may occur, and the new trade-offs that may arise in consequence.

Mechanisms that tie gas exchange and water loss together are often regulated at the respiratory surface (Woods & Smith, 2010). In many amphibians, gas exchange and water loss can be regulated by changes in blood flow (*i.e.* perfusion) to the dermal capillary beds (Burggren & Vitalis, 2005; Riddell et al., 2019) via contraction of smooth muscle around the vasculature, leading to vasoconstriction (Clark & Pyne-Geithman, 2005). In our study, muscle contraction was among the five most significant enriched processes for genes correlated with both changes in resistance to water loss and oxygen uptake. In addition, the module associated with phenotypic change consistent with the trade-off (PC1) was enriched for negative regulation of angiogenesis (*i.e.*, blood vessel growth). Therefore, regulation of the vasculature (such as perfusion, vasoconstriction, and vasoregression) is likely the primary mechanism that shapes the interdependency of resistance to water loss and gas uptake. The analysis also identified potentially novel ways in which regulation of gas uptake may affect resistance to water loss, as in the association between the trade-off and the terms protein degradation and mitochondrion autophagy. The gene with the highest correlation with both traits was *PDK4*, which is involved in the downregulation of aerobic respiration and glucose metabolism (Pettersen et al., 2019). Thus, processes related to the suppression of metabolic rate may result in concomitant reductions in water loss if less blood is delivered to the respiratory surface in the process (for instance, due to lower cardiac output). Furthermore, these top genes were also distributed across many modules, suggesting the trade-off is structured by pathways distributed across the transcriptome. Compensation from alternative pathways may explain the persistence of the trade-off in our focal species as well as many other taxa (Woods and Smith 2010, Riddell et al. 2024). Despite consistency of the trade-off, we also identified periods in which the trade-off was weaker and genes that were associated with independent regulation of both physiological traits.

The independent regulation of gas uptake and skin resistance, although transient, suggests that salamanders have access to phenotypic space outside of the trade-off through phenotypic plasticity, which may allow individuals to produce alternative ecological strategies (Bolnick et al., 2003; Layman et al., 2015; Østman et al., 2014). That is, when selective pressures on the costs of water loss are relaxed (such as due to saturated conditions), metabolic rates may become unconstrained and expressed more freely (Frédérich et al., 2014; Gianoli & Palacio-López, 2009; Pigliucci et al., 1995). The trade-off was not detected in August during the seasonal acclimatization study, which was the most humid month (Supplementary Figure S2). Under these more humid conditions, salamanders could adjust physiological traits more independently due to the lower threat of desiccation. Because we sampled unique individuals at each time point in the seasonal acclimatization study, variation in the trade-off could be due to individual variation rather than phenotypic plasticity. That said, we find this unlikely given the plasticity observed in the laboratory acclimation experiment. Although we did not find an effect of temperature or humidity treatment in the acclimation study, the lack of an effect may have been due to the relatively similar humidity treatments or brevity of the exposure to humidity (∼ 3 h per night). In the gene co-expression network analysis, we identified a module that was related to the independent regulation of gas uptake and skin resistance (*i.e.*, orthogonal to the trade-off axis). This module was enriched with processes mostly related to immune function and unspecific cellular regulatory processes, which likely reflects the underlying processes that drive variation in metabolic rate not explained by changes in skin resistance. In contrast to the network analysis, the correlation analyses were better able to identify putative mechanisms underlying the independent regulation of gas uptake and skin resistance.

Organisms may decouple oxygen uptake from water loss by adjusting processes related to oxygen binding affinity. In our study, genes correlated to the change in oxygen uptake but not skin resistance were enriched for heme binding. Oxygen binding affinity to hemoglobin can be impaired by elevated temperatures due to the exothermic nature of the process (Gangloff & Telemeco, 2018; Pörtner, 2002; Verberk et al., 2016; Weber & Campbell, 2011). Consequently, aquatic and terrestrial organisms generally promote oxygen binding to hemeproteins under hypoxia or temperature stress in acclimation experiments (Chung et al., 2017; Burggren & Wood, 1981; Storz et al., 2010; Tufts et al., 2013). Generally, adaptive increases in oxygen binding affinity are localized to respiratory tissues via changes in temperature, pH, and CO_2_ concentration, and balanced by decreased binding affinity in systemic tissues for oxygen delivery (Webb et al., 2022). In studies on salamanders, some species acclimate by adjusting oxygen binding affinity, whereas others do not (Bonaventura et al., 1977; Maginniss & Booth, 1995; Ultsch, 2012; Weber et al., 1985). Thus, regulating oxygen binding affinity may produce additional trade-offs, particularly if oxygen delivery to systemic tissues is compromised. Similarly, increasing hemoglobin content may promote oxygen carrying capacity of the blood, but at the cost of increased viscosity of the blood (Wood, 1991). Nevertheless, regulation of oxygen binding affinity has the potential to independently adjust gas exchange rates without influencing water loss physiology.

Our analyses also identified putative pathways for the independent regulation of skin resistance to water loss. The expression of tyrosinase (*TYR*) had the strongest correlation with changes in resistance to water loss, while also being uncorrelated to changes in oxygen uptake. *TYR* is the enzyme responsible for oxidizing the amino acid tyrosine, which can then be converted into the various forms of melanin (Sánchez-Ferrer et al., 1995). Several recent studies have indicated that melanin acts as a barrier against water loss in the integument of insects (King & Sinclair, 2015; Rajpurohit et al., 2008; Ramniwas et al., 2013; Välimäki et al., 2015), fungi (Fernandez & Koide, 2013; Jiang et al., 2016), and humans (Elias et al., 2010; Man et al., 2014). The underlying process is related to an acidification effect of melanin in the epidermis which reduces water permeability (Man et al., 2014). The salamanders in our study range from gray to nearly black-pigmented, and being nocturnal, are unlikely to benefit from melanin protection from UV radiation (Nicolaï et al., 2020). Therefore, melanin synthesis may have an important role in water conservation in lungless salamanders. Melanin production, however, likely introduces alternative trade-offs. Its synthesis can be limited by availability of resources such as its amino acid precursor tyrosine or metals such as Cu, Zn, and Fe (McGraw, 2008). Therefore, upregulating the expression of tyrosinase may be costly to other processes (*i.e.*, the Y-model; van Noordwijk & de Jong, 1986). In the absence of these constraints, skin resistance to water loss may be regulated more independently and evolve more freely.

In summary, we identified putative mechanisms that allow organisms to mediate functional conflicts between linked traits. However, while mitigating the trade-off between water loss and oxygen uptake may appear adaptive, alternative resource or functional constraints can emerge. These alternative trade-offs may ultimately preclude lungless salamanders from regulating oxygen uptake and resistance to water loss independently. Because we observed the trade-off persisting through most of the laboratory acclimation experiment and active season, it suggests that escapes from the trade-off may be ephemeral, at least in lungless salamanders. However, even the brief independent regulation of gas uptake and skin resistance may provide ecological advantages if they provide more time to forage and find mates or reduce energetic costs. These results provide new hypotheses on whether the independent regulation of physiological traits allows individuals to outcompete conspecifics or congenerics by mitigating inherent costs of a functional conflict. Such alternative strategies and limitations define the phenotypic space in which natural selection may favor novel trait combinations and allow organisms to occupy new ecological niches.

## Acknowledgements

We thank the North Carolina Wildlife Resource Commission and Nantahala National Forest for permission to conduct our experiments on *P. metcalfi*.

