## Supplementary materials S1-S3 for "Plastic mechanisms for unraveling a universal trade-off between water loss and respiration"

### Effect of laboratory temperature and humidity exposures on the trade-off

The 120 salamanders from the laboratory acclimation experiment were evenly distributed across four experimental groups consisting of temperature and humidity exposures in a 2x2 factorial design. In the *cool* treatment, temperature cycled between 8.5°C and 15°C, while in the *warm* treatment it cycled between 15°C and 22.5°C. The *dry* humidity treatment consisted of a vapor pressure deficit (VPD) of 0.4 kPa, while the *wet* treatment maintained a VPD of 0.2 kPa. We monitored whether VPDs were in the desired levels using three Hygrochron iButtons (Item# DS1923, Maxim Integrated, 160 Rio Robles, San Jose, CA 95134) placed on randomly selected shelves of the incubator each night.

We used linear regression models to investigate the effect of exposure treatment on the trade-off. In model 1, *V*O_2_ was a function of temperature (*cool* or *warm*), humidity (*wet* or *dry*), *r*_i_ , and a three-way interaction between them. Body mass was included as a covariate. Because of high multicollinearity between the predictors (variance inflation factor (VIF) > 10), we also report subsets of this model with only two-way interactions between *r*_i_ and each treatment (models 1a and 1b). Additionally, we investigated the effect of treatment on the change in the trade-off during the experiment. Model 2 was identical to model 1, but with Δ*V*O_2_ and Δ*r*_i_ as dependent and independent variables, respectively. Due to high multicollinearity in model 2 (VIF > 10), we report subsets of this model that only contained two-way interactions between *r*_i_ and each treatment (models 2a and 2b). Type-2 ANOVA outputs for all models are reported in table S1.

Table S1: Type-2 ANOVA outputs for the effects of temperature and humidity exposures on the trade-off between resistance to water loss (*r*_i_) and oxygen uptake. Significant effects would be indicated by significant P-values for any interaction between either a temperature or humidity treatment with final *r*_i_ (model 1) or with Δ*r*_i_ (model 2), which were not observed. Because of high multicollinearity (VIF) in both models, we report subsets of these models having fewer interaction terms.

|  | Predictor | Sum Sq | Df | *F* | *P* | VIF |
| --- | --- | --- | --- | --- | --- | --- |
| Model 1 | Mass | 2628.7 | 1 | 3.74 | 0.056 | 1.19 |
|  | Final *r*_i_ | 1920.0 | 1 | 2.73 | 0.101 | 6.79 |
|  | Humidity | 544.6 | 1 | 0.78 | 0.381 | 259.82 |
|  | Temperature | 4443.6 | 1 | 6.33 | 0.013 | 219.33 |
|  | Final *r*_i_ : Humidity | 385.5 | 1 | 0.55 | 0.460 | 270.64 |
|  | Final *r*_i_ : Temperature | 0.0 | 1 | 0.00 | 0.996 | 227.86 |
|  | Humidity : Temperature | 101.5 | 1 | 0.14 | 0.705 | 295.34 |
|  | Final *r*_i_ : Humidity : Temperature | 352.6 | 1 | 0.50 | 0.480 | 314.38 |
|  | Residuals | 77972.6 | 111 |  |  |  |
| Model 1a | Mass | 2816.3 | 1 | 4.08 | 0.046 | 1.17 |
|  | Final *r*_i_ | 1997.9 | 1 | 2.90 | 0.092 | 3.06 |
|  | Temperature | 4782.2 | 1 | 6.93 | 0.010 | 98.24 |
|  | Final *r*_i_ : Temperature | 6.6 | 1 | 0.01 | 0.922 | 100.91 |
|  | Residuals | 79352.1 | 115 |  |  |  |
| Model 1b | Mass | 2876.5 | 1 | 3.99 | 0.048 | 1.16 |
|  | Final *r*_i_ | 2062.1 | 1 | 2.86 | 0.093 | 2.58 |
|  | Humidity | 548.4 | 1 | 0.76 | 0.385 | 88.13 |
|  | Final *r*_i_ : Humidity | 722.1 | 1 | 1.00 | 0.319 | 91.18 |
|  | Residuals | 82870.4 | 115 |  |  |  |
| Model 2 | Mass | 6600.8 | 1 | 2.11 | 0.149 | 1.06 |
|  | Δ*r*_i_ | 148026.8 | 1 | 47.36 | 0.000 | 6.63 |
|  | Humidity | 73.7 | 1 | 0.02 | 0.878 | 7.93 |
|  | Temperature | 859.9 | 1 | 0.28 | 0.601 | 7.42 |
|  | Δ*r*_i_ : Humidity | 2352.6 | 1 | 0.75 | 0.387 | 13.13 |
|  | Δ*r*_i_ : Temperature | 910.7 | 1 | 0.29 | 0.590 | 12.82 |
|  | Humidity : Temperature | 141.3 | 1 | 0.05 | 0.832 | 10.43 |
|  | Δ*r*_i_ : Humidity : Temperature | 0.0 | 1 | 0.00 | 0.997 | 14.71 |
|  | Residuals | 346940.6 | 111 |  |  |  |
| Model 2a | Mass | 6451.1 | 1 | 2.13 | 0.147 | 1.01 |
|  | Δ*r*_i_ | 151058.1 | 1 | 49.80 | 0.000 | 2.15 |
|  | Humidity | 89.5 | 1 | 0.03 | 0.864 | 3.22 |
|  | Δ*r*_i_ : Humidity | 2526.2 | 1 | 0.83 | 0.363 | 4.36 |
|  | Residuals | 348857.9 | 115 |  |  |  |
| Model 2b | Mass | 6687.5 | 1 | 2.20 | 0.141 | 1.02 |
|  | Δ*r*_i_ | 148253.3 | 1 | 48.76 | 0.000 | 3.06 |
|  | Temperature | 931.4 | 1 | 0.31 | 0.581 | 3.45 |
|  | Δ*r*_i_ : Temperature | 915.5 | 1 | 0.30 | 0.584 | 5.86 |
|  | Residuals | 349626.8 | 115 |  |  |  |

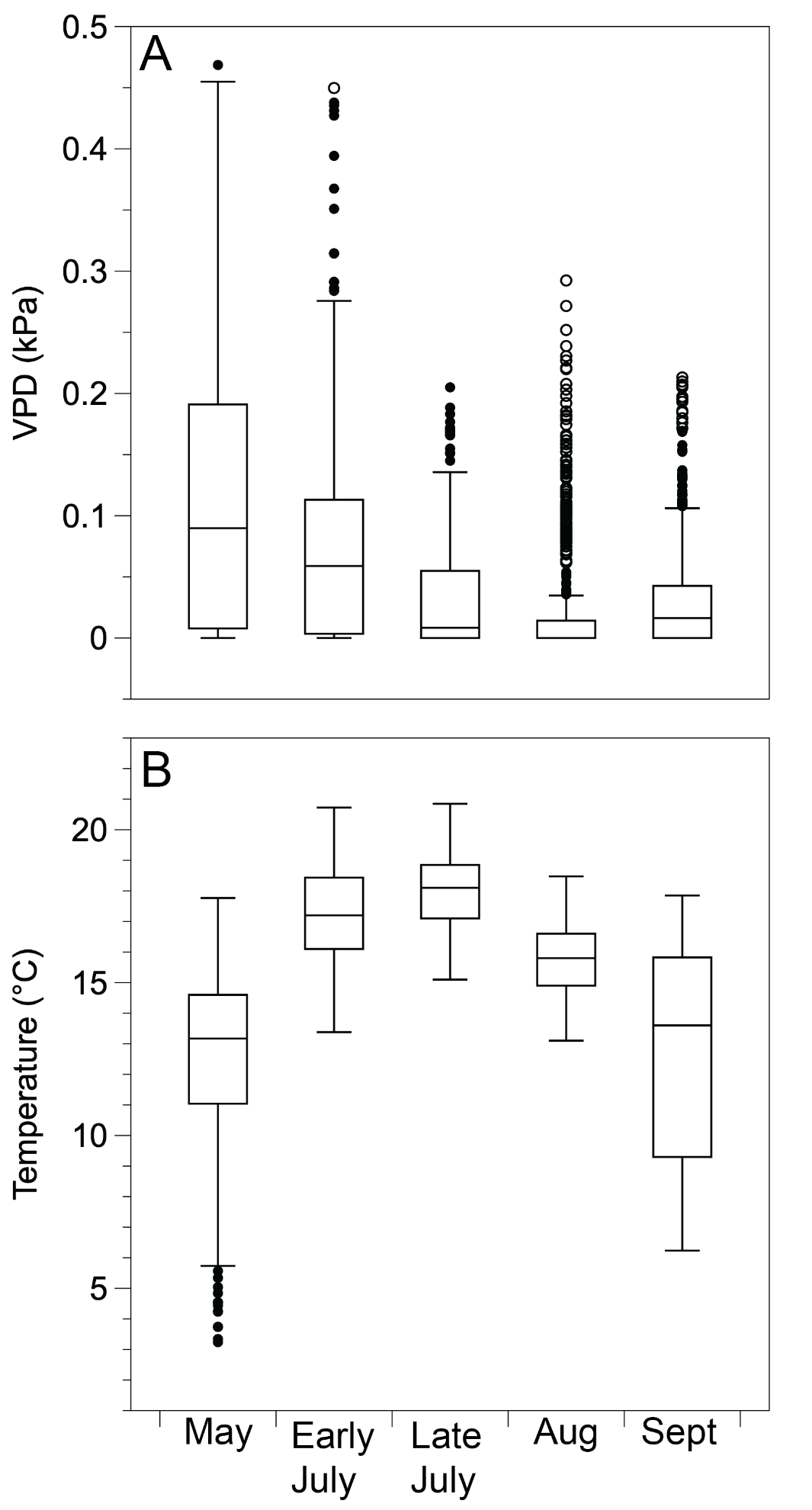

**Figure S2**: Environmental data collected throughout the seasonal acclimatization study from iButtons placed on 15 randomly generated coordinates in the collection sites and in hardwire meshes 1cm above ground. Boxplots are shown for (A) the vapor pressure deficit (VPD) and (B) temperature.

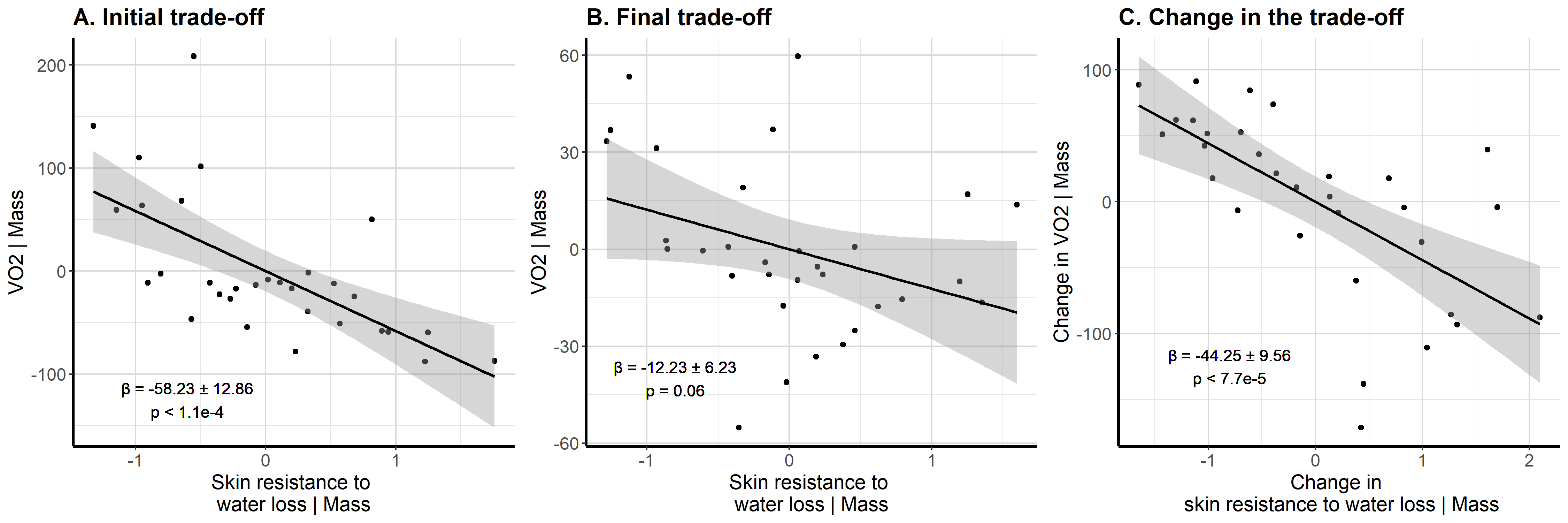

**Figure S3**: For the subset of 32 individuals with gene expression data, (A) the trade-off before acclimation, (B) the trade-off after acclimation, and (C) the trade-off for plasticity in each trait showed qualitatively equivalent results as those obtained from all 120 individuals in the study.
